# The Principal Component Life Trajectory (PCLT): Mapping Sex-Specific Physiological Change Across the Human Lifespan

**DOI:** 10.64898/2026.09.08.750036

**Authors:** Lars Royall, Jordan J. Baechle, Ana Stankovic

## Abstract

Age- and sex-related variation in physiological biomarkers is well documented, yet how coordinated systemic physiology unfolds across life stages at the population level remains unclear. Here, we introduce the Principal Component Life Trajectory (PCLT), reconstructing physiological trajectories via multivariate analysis of 54 blood-based biomarkers from 11,124 participants in the National Health and Nutrition Examination Survey (NHANES). Quantifying year-on-year progression with Euclidean Trajectory Distance (ETD), we identify five distinct phases in an unbiased manner: Early adolescence, Sexual Divergence, Sexual Convergence, Late Adulthood, and Advanced Aging. Trajectories show early, pronounced sexual divergence validated in an ethnically distinct, independent cohort. Chronic disease is associated with PCLT displacement and increased ETD progression, whereas metabolic, behavioural, and psychosocial factors generate graded, phase-specific perturbations. Together, these findings establish PCLT as a reproducible framework for characterizing structured physiological change across the lifespan and for monitoring population-level health dynamics.

## Introduction

Human physiology changes across the lifespan, as reflected in age- and sex-associated variation in circulating biomarkers documented in large population studies (Hägg, 2021; Lau et al., 2019; Lehallier et al., 2019; Moqri et al., 2023; Yang & Kozloski, 2011; Zhernakova et al., 2022). However, how these signals integrate into structured physiological trajectories spanning the full lifespan remains incompletely characterised.

Aging has been defined as the time-related biological and physiological loss of functions necessary for survival and reproduction (Kirkwood & Austad, 2000). However, biological change does not begin abruptly in adulthood nor proceed uniformly across life (Cohen et al., 2023; Lehallier et al., 2019; Oh et al., 2023; Shen et al., 2024; Tian et al., 2023). Different life stages, including development, reproductive maturation, and late life, are associated with distinct patterns of physiological variation (El Khoudary Samar R. et al., 2020; López-Otín, 2013; Veldhuis et al., 2005). Despite broad recognition of nonlinear and sex-specific dynamics, there is limited quantitative reconstruction of how coordinated physiology progresses across these life stages in populations.

Efforts to quantify biological aging have generated composite metrics that estimate biological age or functional decline (Belsky et al., 2015; Bernard et al., 2023; Fuentealba et al., 2025; Horvath, 2013; Levine et al., 2018; Mak et al., 2023). While informative for individual-level risk prediction, such measures provide single age-aligned estimates and do not explicitly reconstruct the physiological progression across life stages in multiple dimensions. As a result, the structured architecture of population-level physiological trajectories remains insufficiently resolved. Prior attempts to characterize age-related patterns using blood biomarkers have often relied on restricted panels such as complete blood counts, offering only a partial representation of systemic physiology and limited insight into sex-specific structure (Pyrkov et al., 2021).

To address this gap, we introduce the Principal Component Life Trajectory framework (PCLT), a population-level multivariate model of lifespan physiology based on principal component analysis of 54 blood-based biomarkers measured in 11,124 participants from the National Health and Nutrition Examination Survey (NHANES). Rather than isolating aging as a discrete phase, this approach reconstructs physiological trajectories from adolescence through advanced age. The framework identifies dominant axes of variation reflecting sexual divergence and temporal progression across the lifespan by quantifying movement along this trajectory using a novel metric termed Euclidean Trajectory Distance (ETD). Using PCLT, we evaluate how chronic disease and distinct internal and external influences on systemic physiology are associated with shifts in trajectory position and progression across life stages.

## Results

### Principal Component Life Trajectory captures age- and sex-associated variation

To map the trajectory of human development and aging, we applied Principal Component Analysis (PCA) to age-sex aggregated physiological profiles, constructed as the mean biomarker values across 11,124 NHANES participants for each sex at each chronological age (12- to 80-year-olds). The objective of this aggregation step was not individual-level prediction but reconstruction of population-level physiological trajectories. This framework, which we term the Principal Component Life Trajectory (PCLT), revealed two dominant and biologically meaningful axes of variation that together accounted for 54% of total variance (Figure 1A, Supplementary Figure 1A). Averaging biomarker values within each age-sex group stabilized inter-individual variance and yielded representative biological profiles across the lifespan (Figure 1A). The first principal component (PC1) primarily captured sex-associated physiological variation, whereas the second (PC2) primarily captured age-associated physiological variation spanning development, maturation, and aging (Figure 1D). To explore the features of these life trajectories, we modelled the PC coordinates against age using segmented linear regression (SLR), (Figure 1B, 1C), which outperformed polynomial models (Supplementary Figure 1C).

**Figure 1.**
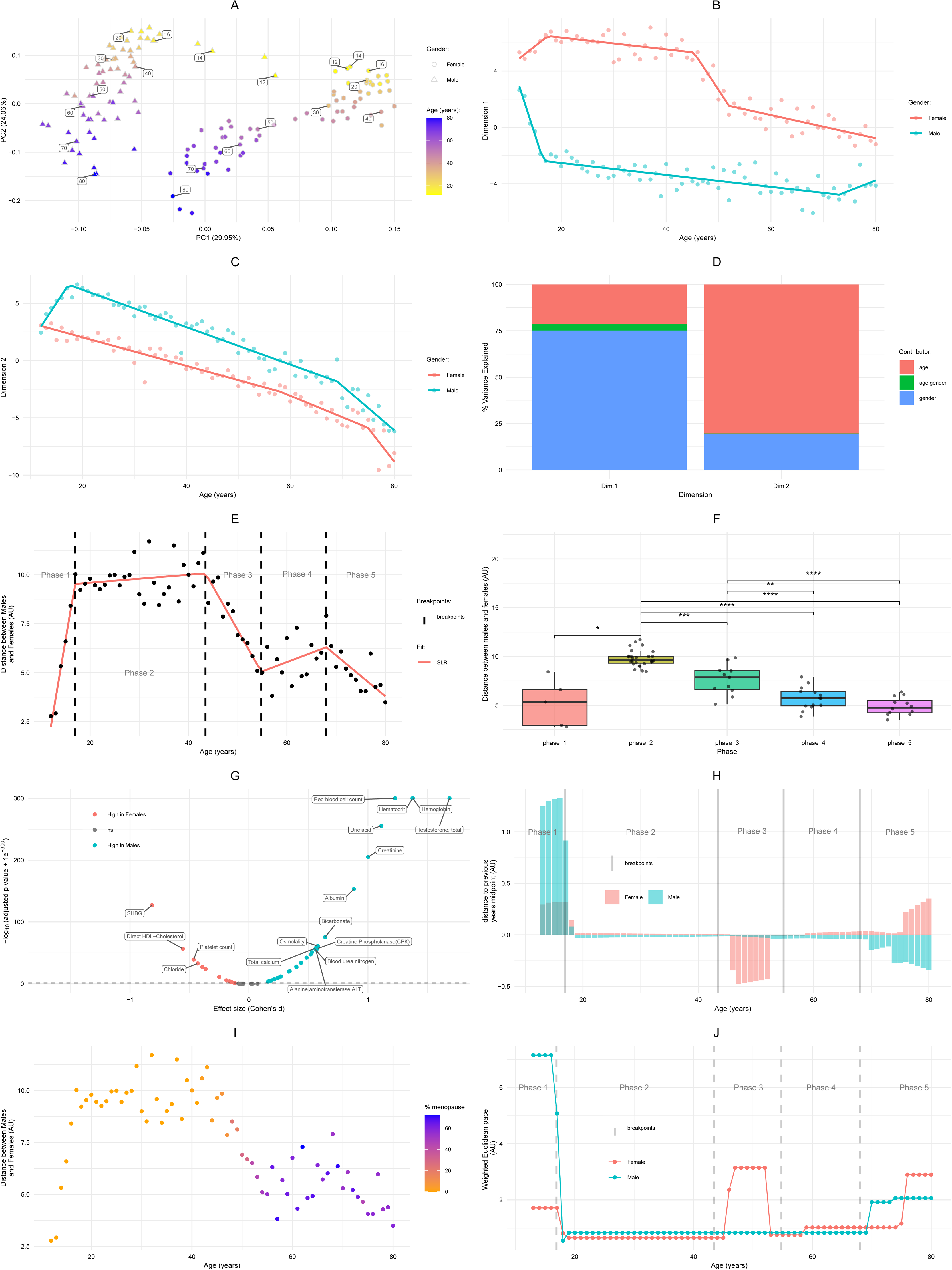
A. Principal Component Analysis (PCA) of mean biomarker profiles from 11,124 NHANES participants aged 12–80, stratified by sex and age. PC1 reflects sex-related variance, while PC2 captures age-related change. B. PC1 values plotted across age show a divergence between males and females beginning in adolescence, peaking in early adulthood, and converging in late life. C. PC2 values plotted across age show a semi-correlated trajectory between the two sexes throughout life, with a divergence during adolescence.. D. Variance in PC1 and PC2 attributed to sex, age, and their interaction. PC1 is primarily sex-driven, while PC2 is primarily age-driven, with some interaction. E. Euclidean distance between male and female trajectories across age, with segmented regression identifying five distinct biological phases: Early adolescence, Sexual Divergence, Sexual Convergence, Late Adulthood, and Advanced Aging. F. Boxplot of Euclidean distance values grouped by identified biological phase. Each phase represents a statistically distinct window of sex divergence. G. Volcano plot showing differential biomarker expression between males and females during the peak divergence window (ages 20–40). Markers such as testosterone and haemoglobin were higher in males; SHBG and HDL cholesterol were higher in females. H. Distance of each sex from the previous year’s intersex centroid, plotted by age and phase. Positive values indicate divergence from the other sex; negative values indicate convergence. Males drive divergence during Sexual Divergence; females drive convergence during Sexual Convergence, consistent with the menopausal transition. I. Euclidean distance between male and female trajectories across age, coloured by the percentage of female participants reporting menopausal status (NHANES variable RHD043). The overlap between peak physiological divergence and rising menopause prevalence is consistent with the menopausal transition being a major contributor to Sexual Convergence. J. Weighted Euclidean Trajectory Distance, plotted by age and sex, highlighting inflection points in the pace of biological change across life. Females show accelerated change from ages 40–60.

In PC1, males and females diverged markedly in adolescence and early adulthood, reflecting the emergence of sex-specific hematic profiles. This divergence peaked during reproductive years and gradually converged in later life; a pattern consistent with age-related reduction in physiological sex differences (Figure 1B).

While PC1 primarily captured sex-associated variation, PC2 represented time-dependent physiological change across development and aging. Plotting PC2 against age revealed a nonlinear trajectory of development and aging in both sexes (Figure 1C). Males exhibit an early divergence from the female trajectory, then advance in parallel before moving toward the females in later life.

To quantify sex-specific separation, we measured Euclidean distance between male and female trajectories across age, which revealed maximum divergence between roughly 20 to 40 years of age (Figure 1E, 1F). We applied SLR to this distance to identify discrete biological stages. This unbiased, data-driven approach partitioned the course of life into 5 distinct stages: Early adolescence, Sexual Divergence, Sexual Convergence, Late Adulthood, and Advanced Aging. Phase boundaries were estimated with narrow confidence intervals for most transitions (e.g., Sexual Divergence onset: age 17.4, 95% CI 13.9–20.9; Sexual Convergence onset: age 45.3, 95% CI 42.9-47.6), supporting the precision of the data-driven segmentation (Supplementary Figure 1D).

We next examined which biomarkers contributed most to this divergence. During the period of maximal dimorphism (ages 20-40), males exhibited higher levels of testosterone, haemoglobin, and creatinine, while females showed elevated SHBG, HDL cholesterol, and platelet counts (Figure 1G). These results confirm that the separation is driven by physiologically meaningful, sex-linked biological changes.

Further exploring how age-specific physiological profiles shifted in PC space revealed that males contributed the most to the separation of the Sexual Divergence, whilst females contributed overwhelmingly to the phase of Sexual Convergence (Figure 1H). This suggests that the menopausal transition contributes substantially to Sexual Convergence, consistent with rising prevalence of reported menopause in this cohort (Figure 1I).

While the PCA coordinates revealed the path of the life trajectory, they did not quantify its pace. To capture the dynamics of biological change, we developed ETD as a variance-weighted metric based on the year-on-year distance moved in PC space, where each dimension’s annual displacement is weighted by the proportion of total variance it explains. ETD thus captures the magnitude of physiological reorganization across the lifespan, with greater weight given to the principal components that account for the most biological variation. This weighting prioritizes trajectory changes occurring along dimensions explaining the largest proportion of physiological variance, ensuring that ETD reflects the dominant axes of systemic change rather than treating all dimensions equally. ETD calculated without variance weighting showed near-identical trajectory patterns (Spearman ρ > 0.99 for both sexes; Supplementary Figure 1E), supporting the robustness of the current formulation. In both sexes, ETD followed a multiphase pattern (Figure 1J): Low, stable rates in early adulthood, a striking midlife acceleration in females (40-60), consistent with the menopausal transition. A secondary small acceleration in both sexes around ages 60-65, and tertiary larger acceleration around 75, suggesting a progressive and systemic decline. These results suggest that population-level physiological change across the lifespan follows a nonlinear, phase-structured trajectory, with the female midlife window representing a major inflection point.

Together, these findings establish PCLT as a framework in which development, maturation, and aging form a single continuum punctuated by inflection points. While PCA captures variance rather than causation, the phase-specific patterns we observe are consistent with established concepts such as developmental programming, epigenetic drift, and inflammaging (Pyrkov et al., 2021; El Khoudary Samar R. et al., 2020; López-Otín, 2013; Veldhuis et al., 2005).

With this approach established, we next investigated how the PCLT framework performs in a demographically and ethnically distinct population. To test whether the PCLT trajectory architecture generalises beyond the NHANES cohort, we performed independent replication in the Korean Health and Nutrition Examination Survey (KNHANES), a demographically and ethnically distinct population with a partially overlapping but reduced biomarker panel (Figure 2B). Remarkably, KNHANES reproduced the sex-structured and nonlinear age trajectory architecture observed in NHANES, supporting cross-population robustness of the PCLT framework (Figure 2A, B, G). Spearman correlation between NHANES and KNHANES PC coordinates was high across all dimensions and sexes (female PC1: ρ = 0.89; female PC2: ρ = 0.96; male PC1: ρ = 0.66; male PC2: ρ = 0.93), confirming that the trajectory architecture is independently replicated rather than merely reproduced. The recovery of similar trajectory architecture despite major differences in ancestry, healthcare systems, recruitment strategies, and biomarker availability suggests that the observed structure reflects fundamental features of human physiology rather than cohort-specific effects.

**Figure 2.**
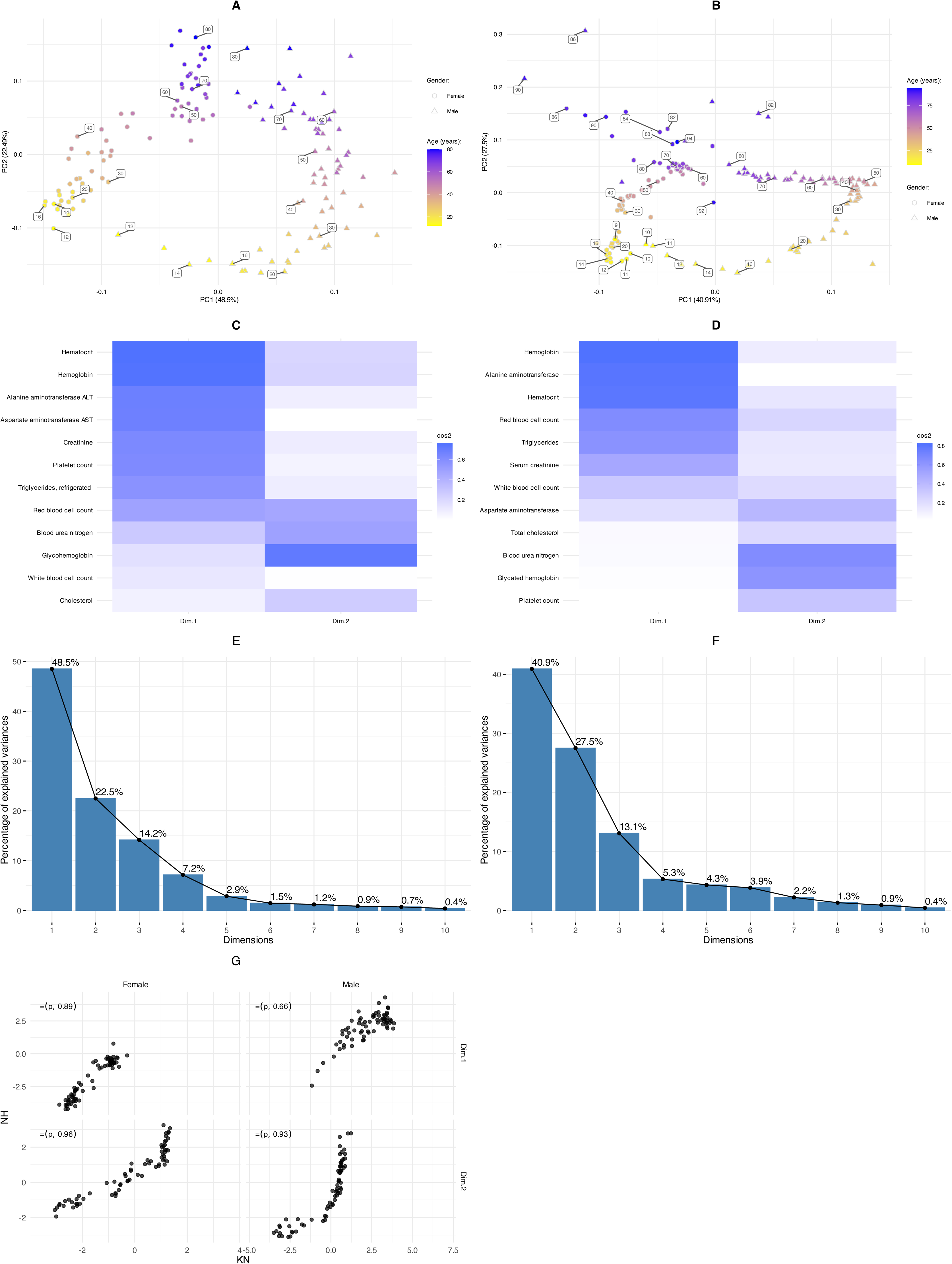
A. Principal Component Analysis (PCA) of mean biomarker profiles from NHANES participants aged 12 years and older, stratified by sex and age, using the minimal biomarker list. B. PCA of mean biomarker profiles from KNHANES participants aged 10 years and older, stratified by sex and age, using the minimal biomarker list. C. Heatmap of cos² values for the NHANES PCA D. Heatmap of cos² values for the KNHANES PCA E. Scree plot showing variance explained by the first 10 dimensions of the NHANES PCA. PC1 and PC2 together account for ∼70.9% of total variance. F. Scree plot showing variance explained by the first 10 dimensions of the KNHANES PCA. PC1 and PC2 together account for ∼68.4% of total variance. G. Spearman’s rank correlation coefficient (ρ) was used to assess the relationship between the PCLT coordinates derived from the KNHANES and NHANES datasets. The analysis demonstrated a strong correlation across all ages, genders, and dimensions, confirming the reproducibility of the PCLT framework across independent cohorts.

### Chronic disease shifts population trajectories toward physiological states characteristic of older ages

Having established the PCLT framework as a map of normative lifespan biology delineating age-related and sex-specific life trajectories, we next applied it to test a central hypothesis in geroscience: if being disease-free was associated with reduced progression along the reconstructed physiological trajectory. To examine this, we compared PCLT derived from the general population (all NHANES participants) with those from a healthy population free of reported chronic disease (Figure 3A). Rather than analysing the smaller, high-variance chronic illness group directly, we subtracted these individuals from the general population to create a more stable baseline for comparison. Artificial individuals were constructed by averaging biomarker values by age, sex, and health status, allowing direct comparisons of PCA positions across populations. Vectors linking each matched age-sex group show that the removal of individuals with chronic illness results in a trajectory shifted toward physiological states characteristic of younger ages, suggesting that the presence of chronic disease in the population is associated with a shift in the average physiological profile toward states characteristic of older ages. This divergence reflects a measurable shift in trajectory position associated with chronic disease burden.

**Figure 3.**
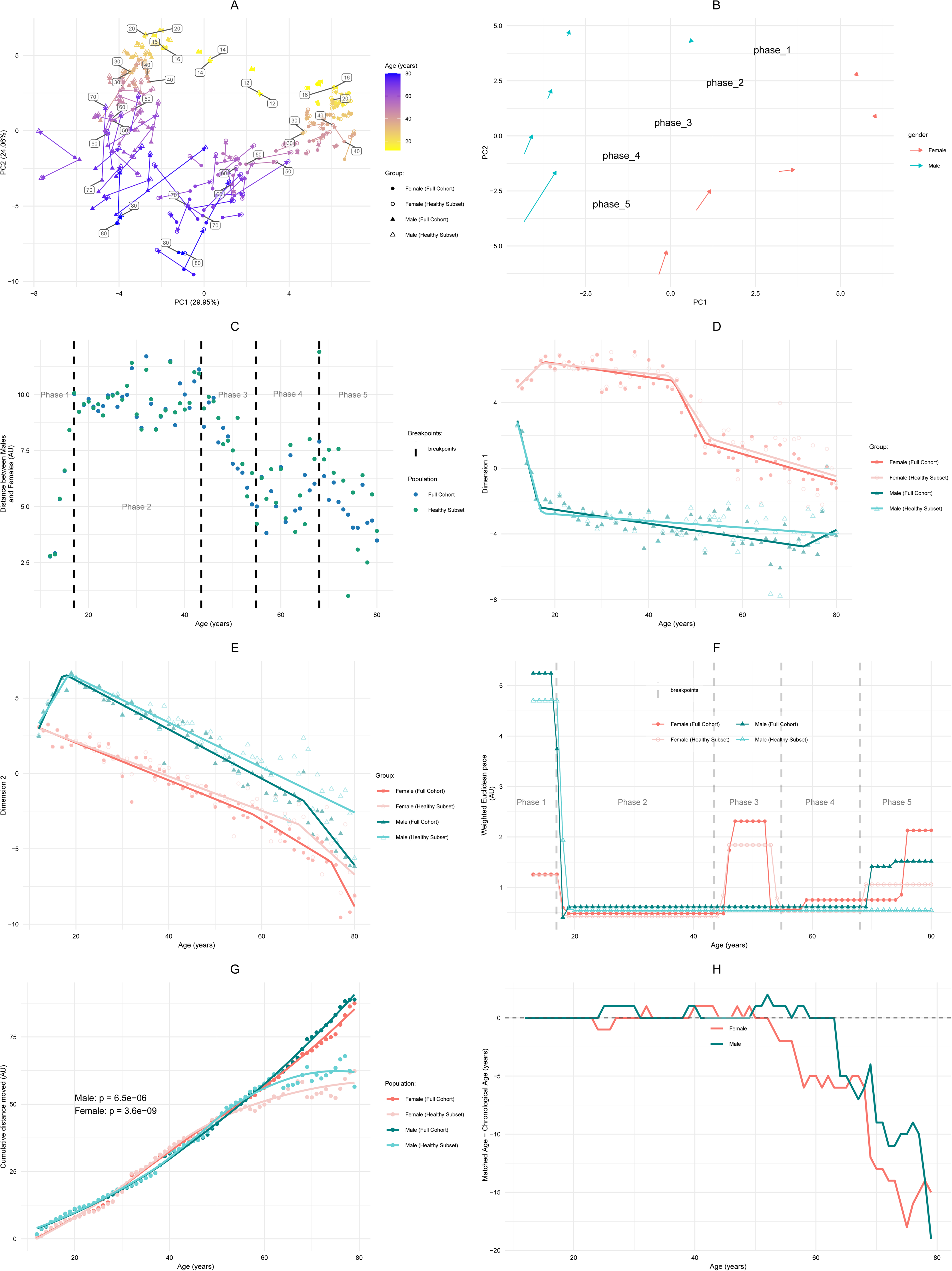
A. PCLT of the general population and healthy subset individuals. The vector arrows indicate the movement of points when individuals previously diagnosed with chronic diseases are removed. B. The mean vector of the change in position between the general population and the age, gender matched healthy subset for each life phase. C. Plot of the distance between age matched genders, for the general population and the healthy cohort. Removal of the chronic disease individuals does not affect the gender distance. D. PC1 values plotted against age show minimal deviation between the full cohort and healthy subset, confirming that chronic disease does not substantially alter sex-specific physiological structure. E. PC2 values plotted against age for all cohorts show the healthy population exhibit an age-related deviation from the general population trend. F. Weighted Euclidean trajectory pace across the four cohorts. Healthy subset males show consistently reduced pace relative to the full male cohort from Phase 2 through Phase 5, with no reversal. Healthy subset females show the same pattern through Phases 1 to 4, but this reverses briefly in early Phase 5, where the healthy subset’s pace exceeds the full cohort’s pace, before the full female cohort’s pace rises sharply again later in Phase 5. In Phase 3, during the menopause transition, healthy females have a lower peak pace compared to the full cohort females. G. Plot of the cumulative distance moved along the PCLT. Females and males of the healthy subset move approximately the same distance as the total population until the approximately the age of 50 and 60 for females and males respectively, where they delineate. H. Plot of the age gap calculated from cumulative distance moved. Both healthy females and males match to younger general population individuals in regards to cumulative distance moved after the age of 50 and 60 for females and males respectively.

To confirm that differences in trajectory were attributable to health status rather than random variation, we validated the healthy population across several dimensions. Individuals without chronic conditions exhibited reduced biomarker variability (Supplementary Figure 2A), lower BMI distributions (Supplementary Figure 2B), and fewer prescription medications (Supplementary Figure 2C). Importantly, sex separation (PC1) was preserved in the healthy population (Supplementary Figure 2D), confirming that observed trajectory shifts reflect disease burden rather than sex-specific clustering.

### Age-dependent divergence peaks in late midlife independent of chronic disease

We next analysed the magnitude and direction of this displacement across lifespan phases. Vectors between matched health states (Figure 3B) show increasing divergence with age, with the largest shifts occurring in Phase 4 (ages 60–70), when trajectory progression increases markedly. Interestingly, sex-specific distance in PCA space remained stable across health groups (Figure 3C), supporting the idea that chronic illness modifies trajectory progression dynamics rather than sexual dimorphism.

To assess how these shifts manifest in trajectory components, we separated the PCA dimensions by group and employed segmented linear regression to model the PCs, which again provided a lower RMSE than polynomial models (Supplementary Figure 2E-F). For the sex dimension (PC1), chronic illness again had minimal impact (Figure 3D). In contrast, the PC2 dimension showed a pronounced elevation in diseased individuals across most ages (Figure 3E), consistent with a shift toward physiological states associated with older ages.

### Chronic disease is associated with accelerated trajectory progression and healthspan compression

To quantify the velocity of these shifts, we calculated ETD on each of the cohorts. Females in the general population exhibited a sharp midlife spike in ETD, which was notably blunted or delayed in healthy females (Figure 3F, Supplementary Figure 2G), suggesting that at least part of the menopausal acceleration is health-state dependent. Males, in contrast, showed a more gradual rise in ETD, with late-life acceleration amplified in the general population group compared to the healthy population.

We next calculated the relative distance between the healthy individuals and the next years’ general population individuals. The complete trajectory for males and females exhibited a significant negative relative decrease between subset and total distance to the next year point. This indicates that the healthy subset exhibits reduced cumulative trajectory progression compared to the general population (Supplementary Figure 2H). By analysing the cumulative distance moved for the healthy and general population, we identified that the populations diverge at around age 50 and 60, for females and males respectively, with the healthy populations exhibiting reduced cumulative trajectory progression compared to the general population (Figure 3G). The timing coincides with the age-associated increase in disease frequency (Gronich et al., 2024; Kuan et al., 2021).

Finally, to express the trajectory divergence in interpretable units, we calculated the Age Gap. For each age in the healthy subset, we identified the closest matching age in the general population by cumulative trajectory progression. The difference between matched age and chronological age quantifies how far ahead along the physiological trajectory the general population is relative to the healthy subset. An Age Gap began emerging at approximately age 50 in females and age 65 in males, expanding rapidly thereafter; by age 79, the healthy subset had a cumulative trajectory progression equivalent to that of a general population individual approximately 16 years younger. The Age Gap thus provides a population-level, trajectory-based summary of the physiological distance between health states across the lifespan.

Together, these findings demonstrate that chronic disease is associated with greater progression along the reconstructed physiological trajectory and with physiological states characteristic of older ages. PCLT thus provides quantitative evidence for healthspan compression and establishes a framework for assessing how biological sex and chronic disease reshape lifespan.

### PCLT detects structured biological perturbations induced by common exposures

Having established that chronic disease produces large-scale displacement in biological trajectories, we next evaluated whether the PCLT framework is sufficiently sensitive to detect more common and graded perturbations. To this end, we examined three distinct factors: obesity (metabolic), smoking (lifestyle), and depression (psychosocial), representing biologically diverse stressors.

Across all three exposures, sex separation in the PC space remained stable (Figure 4B, F, J), indicating that these perturbations, similar to chronic disease described above, do not alter baseline sexual dimorphism. Instead, differences emerged primarily along the age-associated axis of the trajectory dynamics, similar to the impact of chronic disease.

**Figure 4.**
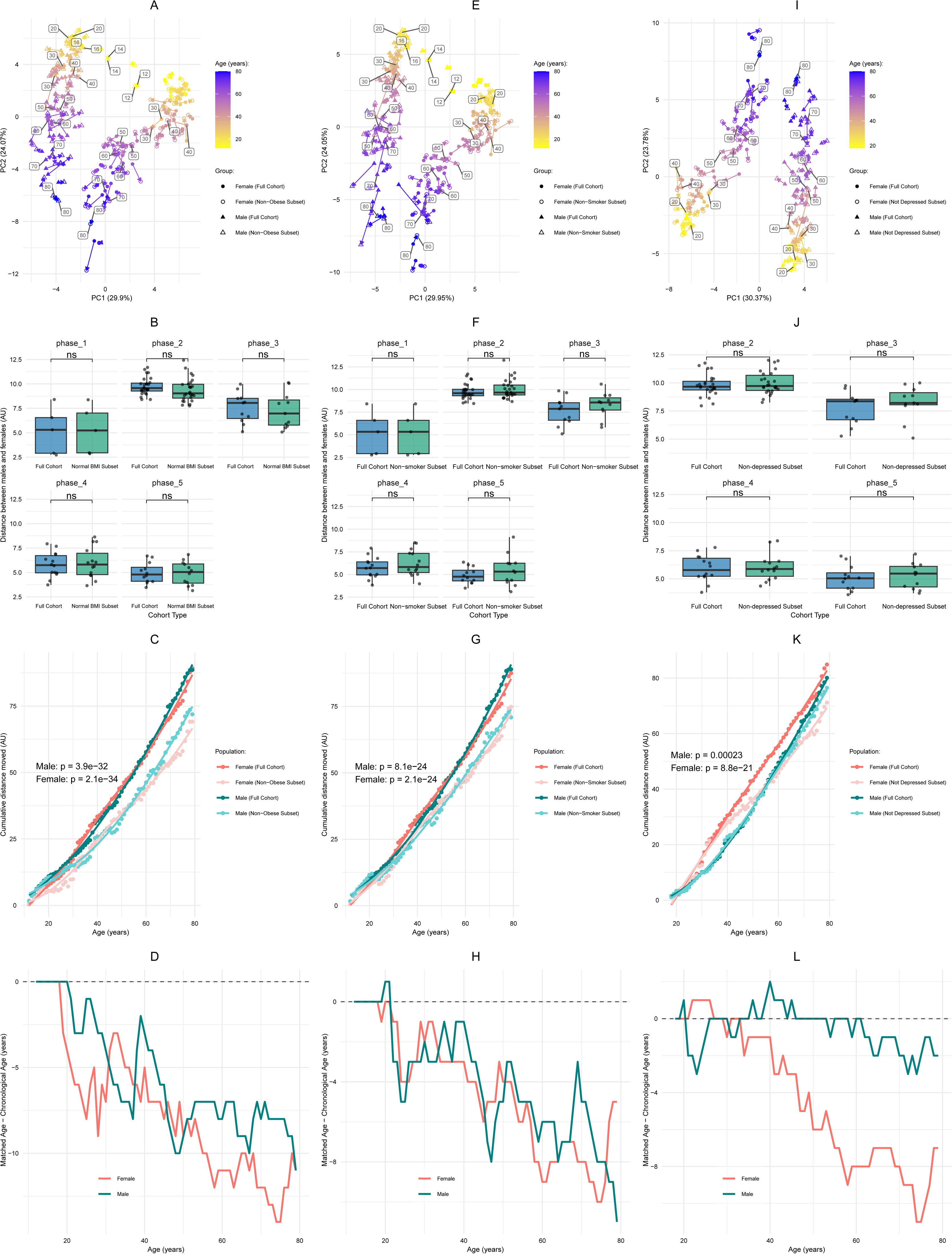
A-D: Obesity is associated with an early shift in physiological trajectory profiles. Non-obese subset population exhibits similar PCA trajectory as the general population (A), and displays no significance deviation in regards to gender difference (B). The cumulative distance moved identifies that the non-obese subset separates quickly from total population during the teenage years (C), which is associated with a large trajectory age gap (D). E-H: Smoking is associated with a trajectory deviation beginning in early adulthood. Non-smoker subset population exhibits similar PCA trajectory as the general population (E), and displays no significance deviation in regards to gender difference (F). non-smoker subset cumulative distance has a more gradual separation from the total and begins in early adulthood (G), which is associated with a moderate trajectory age gap (H). I-L: Depression is associated with sex-specific trajectory effects. Removal of the individuals with depressive symptoms does not affect the PCA (I), and displays no significance deviation in regards to gender difference (J). Non-depressed Females exhibit a strong deviation from the general population, beginning gradually at 30 years old and accelerating at 40 years old. Whereas males have a mild deviation starting later (K). This is associated with a sex-specific trajectory age gap (L).

Removal of obese individuals shifted trajectories toward physiological states characteristic of younger ages, with cumulative divergence emerging in adolescence and widening progressively across life, consistent with compounding metabolic burden and decompensation later in life (Figure 4C-D, Supplementary Figure S3A-C). We next examined smoking as a lifestyle variable and its influence on different life phases. Non-smokers showed a comparable but more gradual reduction in cumulative trajectory distance, with separation beginning in early adulthood and increasing steadily across the lifespan (Figure 4G-H, Supplementary Figure S3D-F).

Finally, we examined the effects of depression. In contrast to smoking and obesity, depression induced subtler and sex-specific effects: females exhibited midlife displacement and earlier cumulative divergence, whereas male effects were delayed and attenuated (Figure 4K-L, Supplementary Figure S3G-I). This is consistent with evidence that female depression risk follows a bimodal pattern, with heightened risk concentrated in adolescence and the menopausal period as critical high-risk windows (Albert, 2015). Interestingly, the age gap identifies a temporal sequence of onset of divergence starting with obesity (Figure 4D), smoking (Figure 4H) and ending with chronic diseases (Figure 3H). This mirrors what is known about the interplay between lifestyle factors and disease risk (Ng et al., 2020).

## Discussion

Human physiology unfolds in a structured and phased manner across the lifespan. Although this has widely been understood to occur, it has remained poorly quantified. Here, we introduce PCLT as a reproducible framework that maps this coordinated architecture of systemic biology and quantifies how populations progress over time. Rather than collapsing biological change into a single scalar estimate, PCLT captures both position within physiological space and the pace of trajectory progression, not only revealing distinct life phases and transitions but also delineating the impact of biological sex and aging.

We demonstrate that chronic disease is associated with measurable displacement and acceleration within this trajectory space. Beyond overt pathology, common metabolic, behavioural, and psychosocial perturbations, including obesity, smoking, and depression, were associated with graded and phase-specific distortions of trajectory dynamics. These analyses indicate that PCLT is sensitive to biologically distinct perturbations while preserving baseline sexual dimorphism.

Most biological age models estimate how old an individual appears biologically by compressing multidimensional physiology into a single scalar value. In contrast, PCLT characterizes age-dependent changes in multidimensional physiological space at the population level, rather than estimating individual biological age. These approaches are complementary, reflecting different levels of biological organization.

The reproducibility of trajectory architecture across geographically and ethnically distinct cohorts, together with reconstruction using tiered biomarker panels, supports the scalability of the framework (Figure 2). Although the analyses rely primarily on cross-sectional and repeated cross-sectional data, which are inherently limited for establishing causality, they demonstrate that coordinated systemic physiology can be mapped and compared across populations in a structured and interpretable manner. As a novel metric of population-level trajectory progression, ETD complements existing longitudinal measures such as DunedinPACE by operating at the population rather than individual level.

Taken together, our findings define PCLT as a robust and reproducible framework that captures the phased architecture of human systemic physiology across the lifespan, quantifies trajectory displacement and acceleration induced by disease and common exposures, and enables dynamic monitoring of population-level health.

Several limitations of the present study should be noted. First, PCLT is reconstructed from cross-sectional and repeated cross-sectional data, meaning that the trajectories represent population-level differences across age rather than longitudinal change within individuals; causal inferences cannot be drawn from this design. Longitudinal cohorts capable of directly validating these trajectories generally involve trade-offs in sample size, biomarker breadth, or coverage of the full age range examined here. The recovery of a smooth, phase-structured trajectory from independently estimated age-specific population profiles, together with the reproduction of its broad architecture in the independent KNHANES cohort, is therefore notable, but does not substitute for longitudinal validation. Future studies applying PCLT to repeated measurements within individuals will be required to determine the extent to which the population-level trajectory identified here reflects within-person physiological change. Second, PCA identifies dominant axes of variance rather than biological mechanisms, and the labelling of PC1 as sex-associated and PC2 as age-associated reflects empirical observation rather than causal structure. Third, all disease and exposure analyses are observational, and the associations reported should not be interpreted as evidence that disease or lifestyle factors directly cause trajectory displacement. Fourth, ETD is a novel metric that has not yet been validated against established longitudinal measures. Future work linking ETD-derived phase transitions to longitudinal cohort data will be necessary to establish its predictive validity. Additionally, breakpoint precision varied across dimensions; the female PC2 trajectory showed a wider confidence interval for the first breakpoint (95% CI: 45.9–70.1 years), reflecting greater variability in the age-associated trajectory dimension for females. Fifth, the Age Gap is a population-level trajectory summary and does not constitute an individual biological age estimate. Finally, biomarker panel composition differs between NHANES and KNHANES, which limits direct comparability of absolute trajectory positions across cohorts.

## Acknowledgments

We thank Prof. Graham Pawelec for critically reviewing the manuscript and for his contributions to the conceptual development of this study. We also thank the National Health and Nutrition Examination Survey (NHANES) and Korean Health and Nutrition Examination Survey (KNHANES) staff and participants, whose data collection efforts made this work possible.

## Ethics Approval and Consent to Participate

This study uses publicly available, de identified survey data from NHANES and KNHANES. No additional ethics approval was required for this secondary analysis.

## Data Availability

NHANES data are publicly available from the Centers for Disease Control and Prevention at https://wwwn.cdc.gov/nchs/nhanes/. KNHANES data are publicly available from the Korea Disease Control and Prevention Agency at https://knhanes.kdca.go.kr/knhanes/main.do.

## Code Availability

Code supporting the findings of this study has been deposited in a public GitHub repository, which will be made publicly available upon submission of the manuscript for peer review. Prior to this, the code is available upon reasonable request.

## Author Contributions

L.R., J.B. and A.S. conceptualized the study. L.R. performed the data analysis and drafted the manuscript. J.B. and A.S. supervised the study and contributed to data interpretation. All authors reviewed, edited, and approved the final manuscript.

---

Figure S1 A. Scree plot showing variance explained by the first 10 PCA dimensions. PC1 and PC2 together account for ∼54% of total variance.

B. Heatmap of cos² values (PCA contribution strength) for individual biomarkers across PC1 and PC2. PC1 is driven by sex-linked markers (testosterone, haemoglobin, haematocrit, creatinine, SHBG), while PC2 loads primarily on markers related to micronutrient status, including folate metabolites and vitamin D.

C. Comparison of segmented vs. polynomial regression models for PC1 and PC2 trajectories. Segmented models consistently showed lower RMSE, especially in PC2 and female data.

D. Breakpoint estimates with 95% confidence intervals for all phase boundaries, derived from the segmented linear regression models fitted to PC1 and PC2 trajectories for females and males. Confidence intervals reflect the standard errors reported by the segmented R package. E. Normalized Euclidean Trajectory Distance (ETD) plotted by age and sex for four alternative ETD formulations: weighted PC1+PC2 (current method), unweighted PC1+PC2, weighted PC1–PC5, and unweighted PC1–PC5. All variants are normalized to their group mean for direct comparison. The female midlife peak and late-life acceleration are consistent across formulations, supporting the robustness of the current approach.

Figure S2 A. Variance of chronic disease population normalised to the variance of the healthy subset.

B. Comparison of the number of prescription medications taken by the chronic disease and healthy populations

C. Comparison of the Body Mass Index (BMI) of the chronic disease and healthy populations

D. Comparison of the gender distance between the two populations reveal no statistically significant change in any of the phases.

E., F. Comparison of segmented vs. polynomial regression models performance for all PCs, gender and populations

G. Comparison of the distance moved per year of each population for the different phases of the PCLT.

H. Boxplots of the relative distance to the next year point indicate that both the healthy subset populations move less through the PCLT, which mainly occurs in phases 4 and 5 of the PCLT

Figure S3 A, D, G. The subset’s relative distance to the next year’s point was calculated in relation to the total population’s distance. Boxplots represent the relative distance for the whole PCLT as well as for each phase. Obesity has a significant effect on relative distance for both genders (A), as does smoking (D). Whereas, depression effect is seen in males, but females particularly affecting phases 2 and 3 (G). All subsets that have the conditions removed have a negative relative change suggesting that the conditions serve to push the individuals further along the PCLT.

B, E, H. A 4 Degree polynomial modelled the relationship between age and relative distance for both genders for each of the three conditions.

C. The rate of obesity against age for this dataset.

F. Plot of the rate of chronic disease in the dataset, suggests that the separation in the 3 conditions is not related to a peak in chronic diseases.

I. The rate of depression again age for this dataset

## Methods

The primary study population was created from the National Health and Nutrition Examination Survey (NHANES) from 2013 to 2016. A detailed description of the survey and its protocols can be found in the NHANES manual (Centers for Disease Control and Prevention/National Center for Health Statistics. Nhanes survey methods and analytic guidelines. 2023. https://wwwn.cdc.gov/nchs/nhanes/analyticguidelines.aspx). Raw data files were downloaded from the NHANES website. Participants under the age of 12 were excluded from the study cohort. Variables with more than 20% missing values were dropped; then, all participants remaining that had missing values were excluded. In total, 11,124 remaining participants were included in our study cohort. NHANES survey weights were not applied in the construction of age-sex mean biomarker profiles. Because PCLT is designed as a descriptive reconstruction of population-level physiological structure rather than an estimate of nationally representative population parameters, unweighted age-sex means were used to define the trajectory. This choice prioritizes the internal geometry of the biomarker space over population-level prevalence estimation; applying survey weights would be necessary for inferential population estimates but does not alter the structural relationships that PCLT reconstructs.

| Category | Biomarkers |
| --- | --- |
| Blood Cell Counts | Basophils number (1000 cells/uL), Eosinophils number (1000 cells/uL), Lymphocyte number (1000 cells/uL), Monocyte number (1000 cells/uL), Segmented neutrophils number (1000 cell/uL), Haematocrit (%), Haemoglobin (g/dL), Mean cell haemoglobin concentration (g/dL), Mean cell haemoglobin (pg), Mean cell volume (fL), Mean platelet volume (fL), Platelet count (1000 cells/uL), Red blood cell count (million cells/uL), Red cell distribution width (%), White blood cell count (1000 cells/uL) |
| Lipids | High density lipoprotein-Cholesterol (HDL, mmol/L), Cholesterol (mmol/L), Triglycerides (mmol/L) |
| Blood Glucose | Glycosylated haemoglobin (HbA1c, %), Glucose (mmol/L) |
| **Kidney & Liver Function** | Albumin (g/L), Blood Urea Nitrogen (BUN, mmol/L), Creatinine (umol/L), Globulin (g/L), Total Bilirubin (umol/L), Total protein (g/L), Uric acid (umol/L), Alkaline Phosphatase (ALP, IU/L), Aspartate Aminotransferase (AST, U/L), Alanine Aminotransferase (ALT, U/L), Gamma Glutamyl Transferase (GGT, IU/L), Lactate dehydrogenase (U/L) |
| **Hormones** | Oestradiol (pg/mL), Sex Hormone Binding Globulin (SHBG, nmol/L), Testosterone (ng/dL) |
| **Vitamins & Minerals** | 25-hydroxyvitamin D2 (25OHD2, nmol/L), 25-hydroxyvitamin D3 (25OHD3, nmol/L), epi-25OHD3 (nmol/L), Red blood cell folate (nmol/L), Serum total folate (nmol/L), Folic acid (nmol/L), 5-Formyl-tetrahydrofolate (nmol/L), Tetrahydrofolate (nmol/L), 5,10-Methenyl-tetrahydrofolate (nmol/L), Mefox oxidation product (nmol/L), Calcium (mmol/L), Iron (umol/dL), Phosphorus (mmol/L), Bicarbonate (mmol/L), Chloride (mmol/L), Potassium (mmol/L), Sodium (mmol/L), Osmolality (mmol/Kg) |
| **Muscle enzyme** | Creatine Phosphokinase (CPK, IU/L) |

The validation dataset was created from Korean Health and Nutrition Examination Survey (KNHANES) data from 1998 to 2020. Information about the study and its data can be found at https://knhanes.kdca.go.kr/knhanes/main.do. Due to differences in age range of tested biomarkers, participants aged between 9 and 94 were kept. Variables associated with blood biomarkers were kept, excluding those relating to heavy metals. Rows were removed if they were variables were completely missing, then columns dropped if they had more than 50% missingness. Finally, any remaining rows with missing values were removed. This resulted in a dataset of 74,500 participants and 12 blood biomarkers. A dataset with matching variables was created from the previously described NHANES dataset.

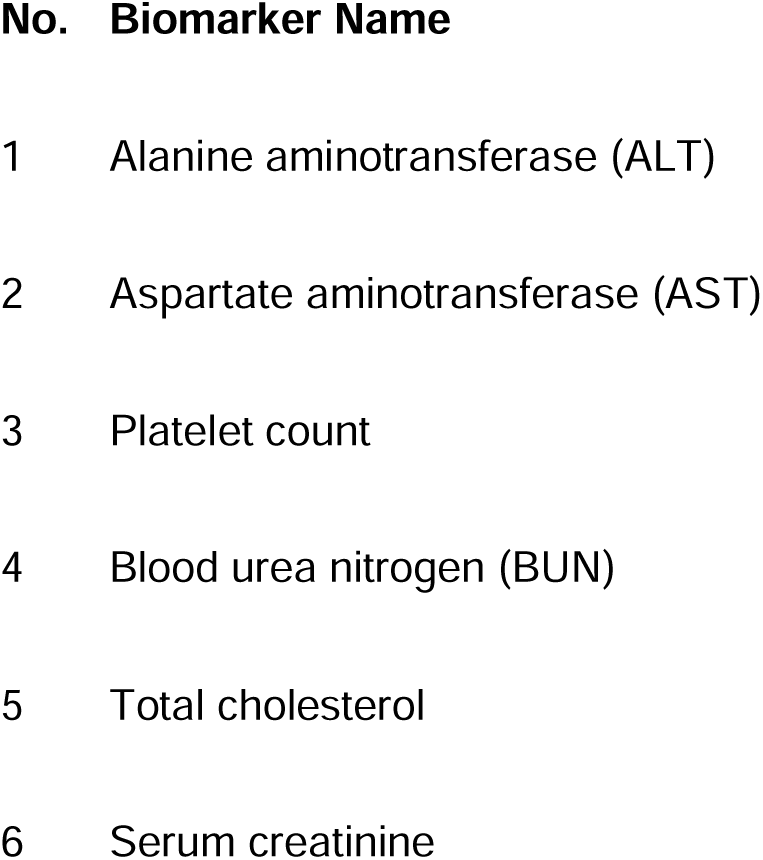
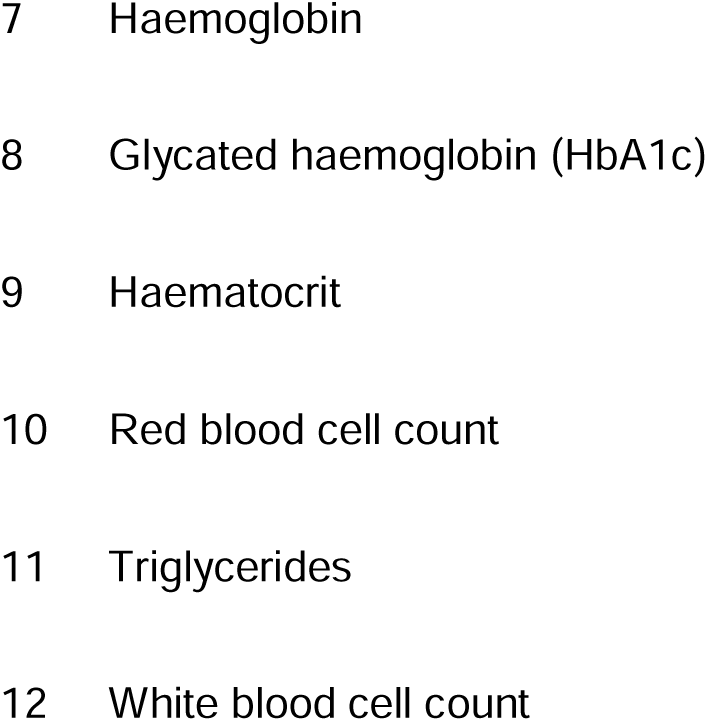

A mean value for each biomarker was calculated for each age and gender combination in the dataset. After excluding variables with a variance of 0, PCA was performed on the age-gender mean data set using the r package ‘stats’ (https://stat.ethz.ch/R-manual/R-devel/library/stats/html/00Index.html). Scree plot and Cos2 for the variables was extracted from the pca using the factoextra r package.

Models were made using between 1 and 10 degrees or breakpoints for polynomial regression and segmented linear regression respectively. Data was split into train (2/3) and test (1/3), the model created and the Root Mean Square Error (RMSE) calculated; this was performed 3 times iteratively on a different 1/3 test data. Models that failed to be created 2 out of 3 times were dropped. The mean RMSE for each model was taken and used to determine the best performing model and its parameter (no. degrees/breakpoints). Finally, a model was created on the full data to capture the entire variance of the data. If Segmented Linear Regression was unable to fit to the full data, the number of breakpoints was reduced by 1. Contributions to the model were calculated by performing Anova analysis on a gender pooled model, extracting the Sum Squares of each factor and representing it as a percentage of the total explained variance.

The distance between males and females was calculated by extracting the first- and second-dimension coordinates of each sex-specific average age from the PCA and using the formula:

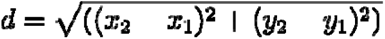

Phases were calculated using the breakpoints and their locations extracted from a model created using the previously defined method of model fitting. Statistical tests were performed between male vs female biomarkers ages between 20 to 40, where it was observed that there was maximal distance between the sexes. Student t.test were performed and the resulting p values were corrected for multiple comparisons (Bonferroni method). The effect size of the difference was calculated using Cohen’s d. 1e-300 was added to all p values to allow for visualization during plotting.

To quantify the directional migration of each sex within PC space, a centroid was calculated at each age t as the midpoint between the male and female PC coordinates. For each sex, the distance from the previous year’s centroid to the current year’s male and female points was computed, and compared to the distance from the current year’s centroid to the current year’s points. The annual change in this distance reflects how much closer to or further from the intersex centroid each sex has moved in a given year: positive values indicate movement away from the other sex (divergence), and negative values indicate movement toward the other sex (convergence). This is expressed in the following equations.

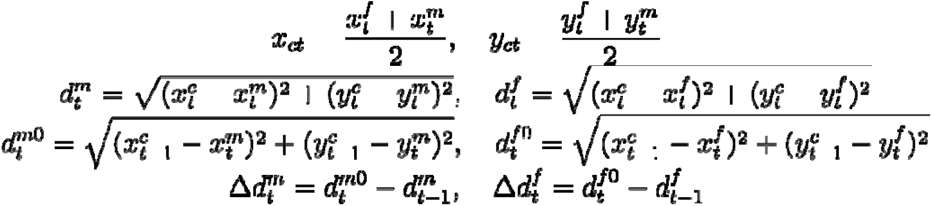

Menopause data was extracted from the NHANES questionnaire variable “RHD043 - Reason not having regular periods”.

The annual Euclidean Trajectory Distance (ETD) was calculated for each sex and population as the variance-weighted year-on-year displacement in PC space. Specifically, ETD is defined as the square root of the sum of squared annual displacements along each PC dimension, weighted by the proportion of total variance explained by that dimension, which is expressed as this equation. Here, w_x and w_y are the proportions of total variance explained by PC1 and PC2 respectively, x_t and y_t are the PC1 and PC2 coordinates at age t, and x_(t-1) and y_(t-1) are the coordinates at age t-1. Variance proportions were derived from the PCA of the full cohort and applied consistently across all subgroup analyses, such that trajectory distances are expressed in a common reference frame.

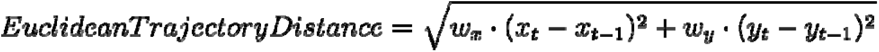

Correlation between NHANES and KNHANES coordinates for the different dimensions and genders was performed with Spearman’s rank correlation (Spearman’s ρ) using the ggpubr library.

Chronic disease status was ascertained from Diabetes and Medical Conditions questionnaire answers. Participants were categorized as having a chronic disease if they answered they had at least one of the following diseases: diabetes, arthritis, congestive heart failure, coronary heart disease, angina pectoris, heart attack, stroke, any liver condition, jaundice. Mean values for each biomarker were again calculated for each age, gender, and chronic disease status combination, and the PCA plotted. Distance between chronic disease participants and non-chronic disease participants used the previously described distance calculation for male and female distance. Similarly, the weighted Euclidean trajectory distance was calculated using the above-described method for both chronic and nonchronic participants.

To compare biomarker variance for individuals with and without chronic disease, the variance of each biomarker among participants older than 50 years was calculated for each group. The variance of the chronic disease group was normalised to the variance of the non-chronic group, yielding a relative variance measure. The number of prescription medications taken by the chronic and non-chronic groups was extracted from the NHANES variable “RXDCOUNT: The number of prescription medicines taken”. Body mass index (BMI) of individuals was calculated using the standard clinical formula of:

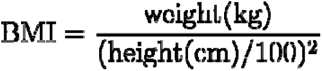

To subset the total population for the three conditions (Obesity, Smoking and Depression) a population with the condition was defined and then removed from the total population. Obesity was defined as having a BMI of greater than 30. Smoking status was defined using the variable “SMQ020 - Smoked at least 100 cigarettes in life”. In order to have a sufficiently high n, depression was defined using “Mental Health - Depression Screener”, which incorporates the DSM-IV depression diagnostic criteria. A score of greater than 5 indicated whether an individual has mild depressive symptoms. For all three conditions, mean biomarker profiles were aggregated by age, sex, and subset status, and PC coordinates for each subset were obtained by projecting the scaled subset data onto the full cohort PCA space, ensuring that all subset trajectories are expressed in a common reference frame consistent with the chronic disease analyses.

Distance to the next year’s point for each gender was calculated for the total and subset populations using the previously described formula for Euclidean distance. The relative change in distance between the total and the subset was calculated using the below formula. To mitigate the influence of extreme outliers, relative distance was Winsorized at the 2.5th and 97.5th percentiles. Two-sided Wilcox signed-rank tests were performed on relative distance for each phase and gender separately. This non-parametric test evaluated the null hypothesis that the median of the distribution was zero.

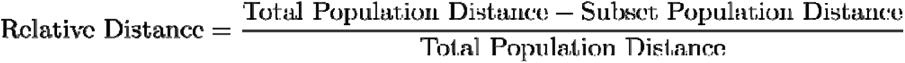

Cumulative distance was calculated separately for the full cohort and each subset population as the running sum of each population’s own year-on-year ETD values from age 12 onward. Because subset individuals are projected onto the full cohort PCA space, cumulative distances are directly comparable across populations. Divergence between the full cohort and subset cumulative distance curves therefore reflects genuine differences in the pace of biological change rather than differences in the coordinate system. The age at which subset and full cohort curves begin to separate identifies the life stage at which the condition of interest becomes associated with increased progression along the reconstructed physiological trajectory. To test whether trajectories differed between the total and subset populations, we fitted cubic polynomial regression models including an interaction between age and population type. Models were fitted separately for males and females. The significance of population differences was assessed using likelihood ratio tests comparing models with and without the age × population interaction.

The Age Gap between the general population and the healthy subset was computed for each gender by first finding the matched age which has the most similar accumulated distance. Age Gap was then calculated as the Matched Age - Chronological Age.

During preparation of this manuscript, the authors used Claude (Anthropic) to check grammar, spelling, and phrasing in text the authors had already written. The authors reviewed, edited, and approved all AI assisted content and take full responsibility for the accuracy and integrity of the manuscript.

