## Supplementary Figure 1 for "The Principal Component Life Trajectory (PCLT): Mapping Sex-Specific Physiological Change Across the Human Lifespan"

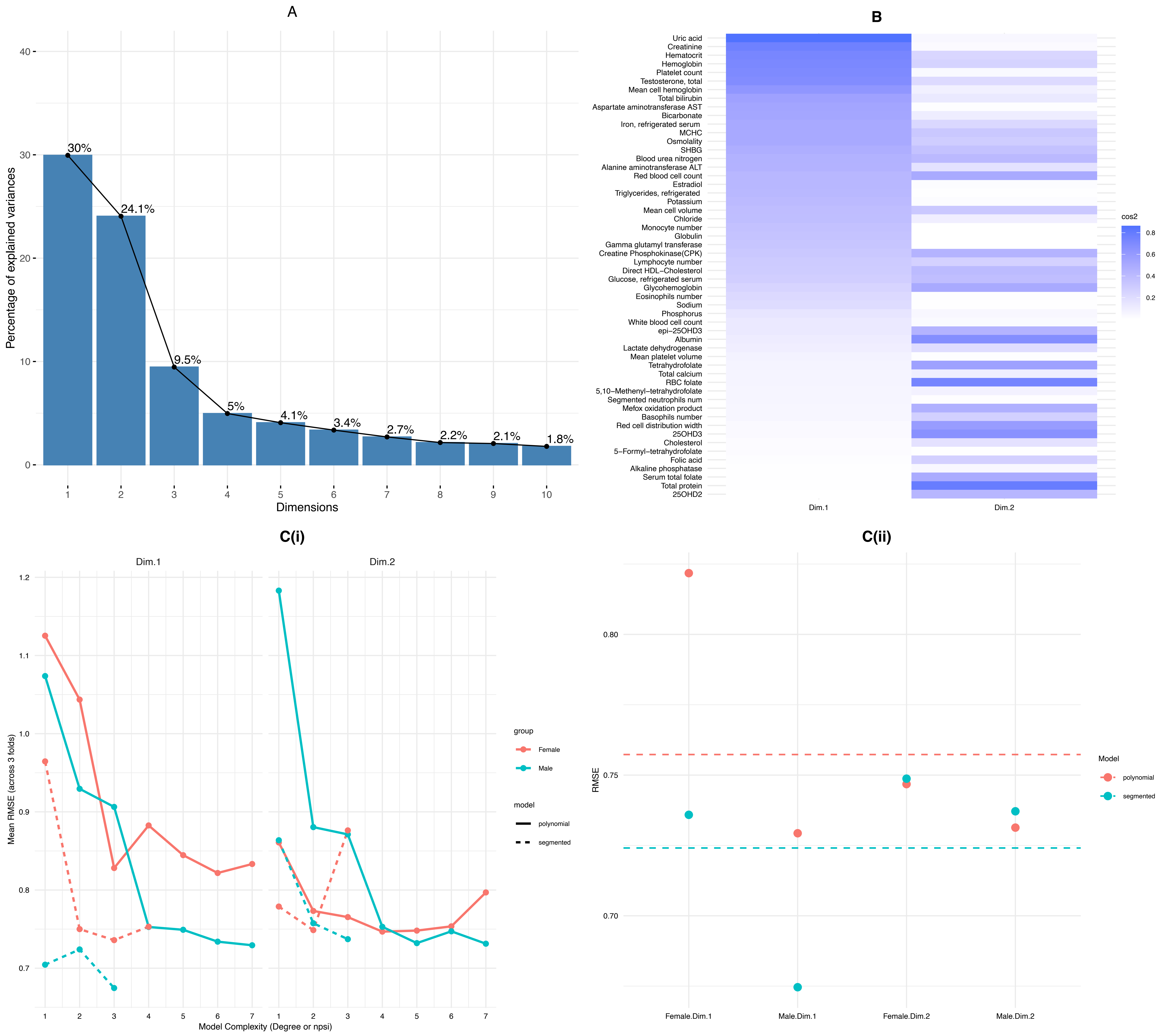


## D

| Sex | PC Dimension | Breakpoint | Age Estimate | SE | Lower CI | Upper CI |
|---|---|---|---|---|---|---|
| Female | Dim.1 | psi1.age | 17.4 | 1.8 | 13.9 | 20.9 |
| Female | Dim.1 | psi2.age | 45.3 | 1.2 | 42.9 | 47.6 |
| Female | Dim.1 | psi3.age | 52 | 1.4 | 49.3 | 54.7 |
| Female | Dim.2 | psi1.age | 58 | 6.2 | 45.9 | 70.1 |
| Female | Dim.2 | psi2.age | 74.9 | 1.6 | 71.9 | 78 |
| Male | Dim.1 | psi1.age | 16.6 | 0.6 | 15.4 | 17.8 |
| Male | Dim.1 | psi2.age | 73 | 2.6 | 68 | 78 |
| Male | Dim.2 | psi1.age | 17.3 | 0.7 | 16 | 18.7 |
| Male | Dim.2 | psi2.age | 69 | 1.9 | 65.3 | 72.7 |

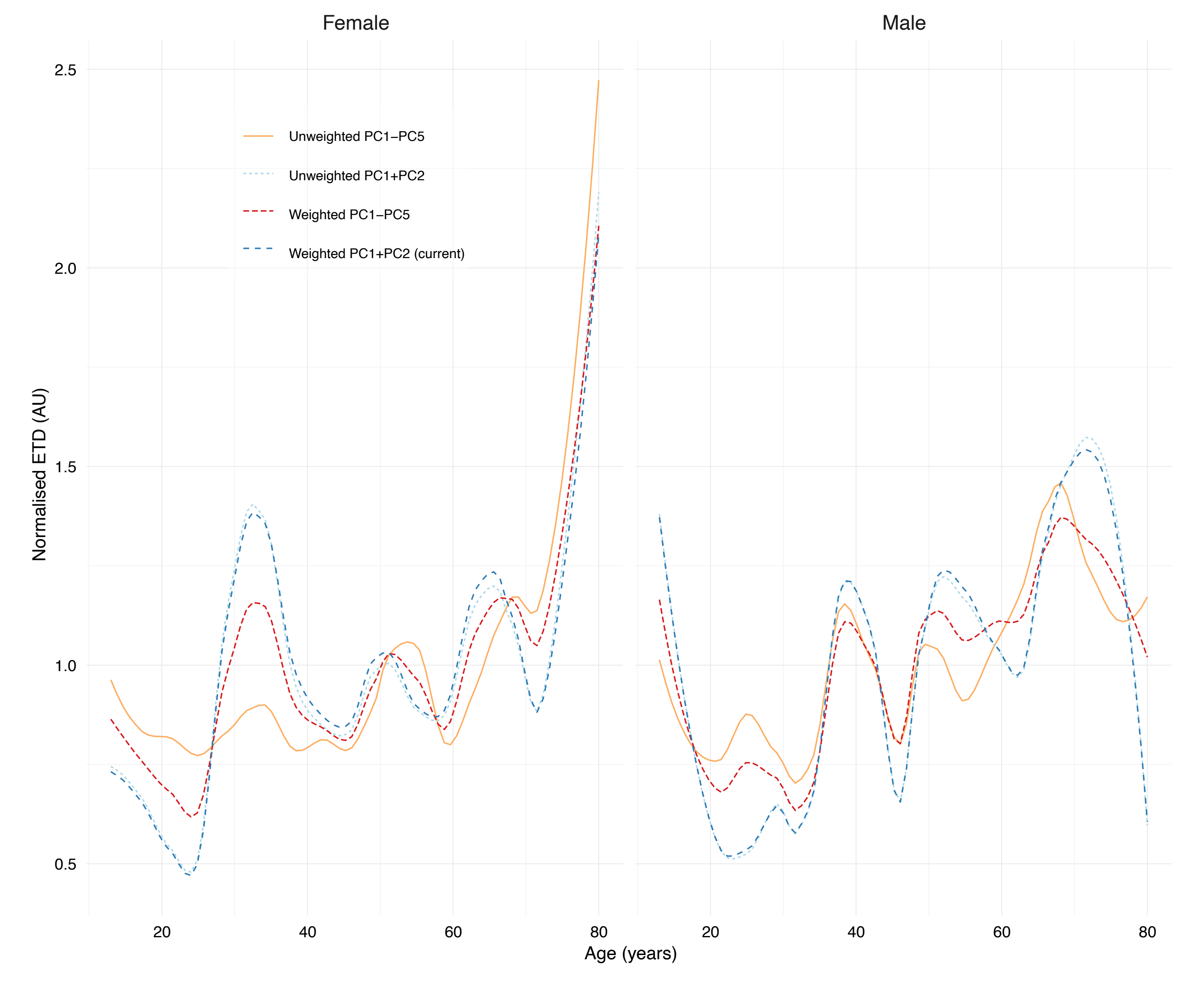
