## Supplementary figures and images for "The Principal Component Life Trajectory (PCLT): Mapping Sex-Specific Physiological Change Across the Human Lifespan"

### Supplementary Figure 2

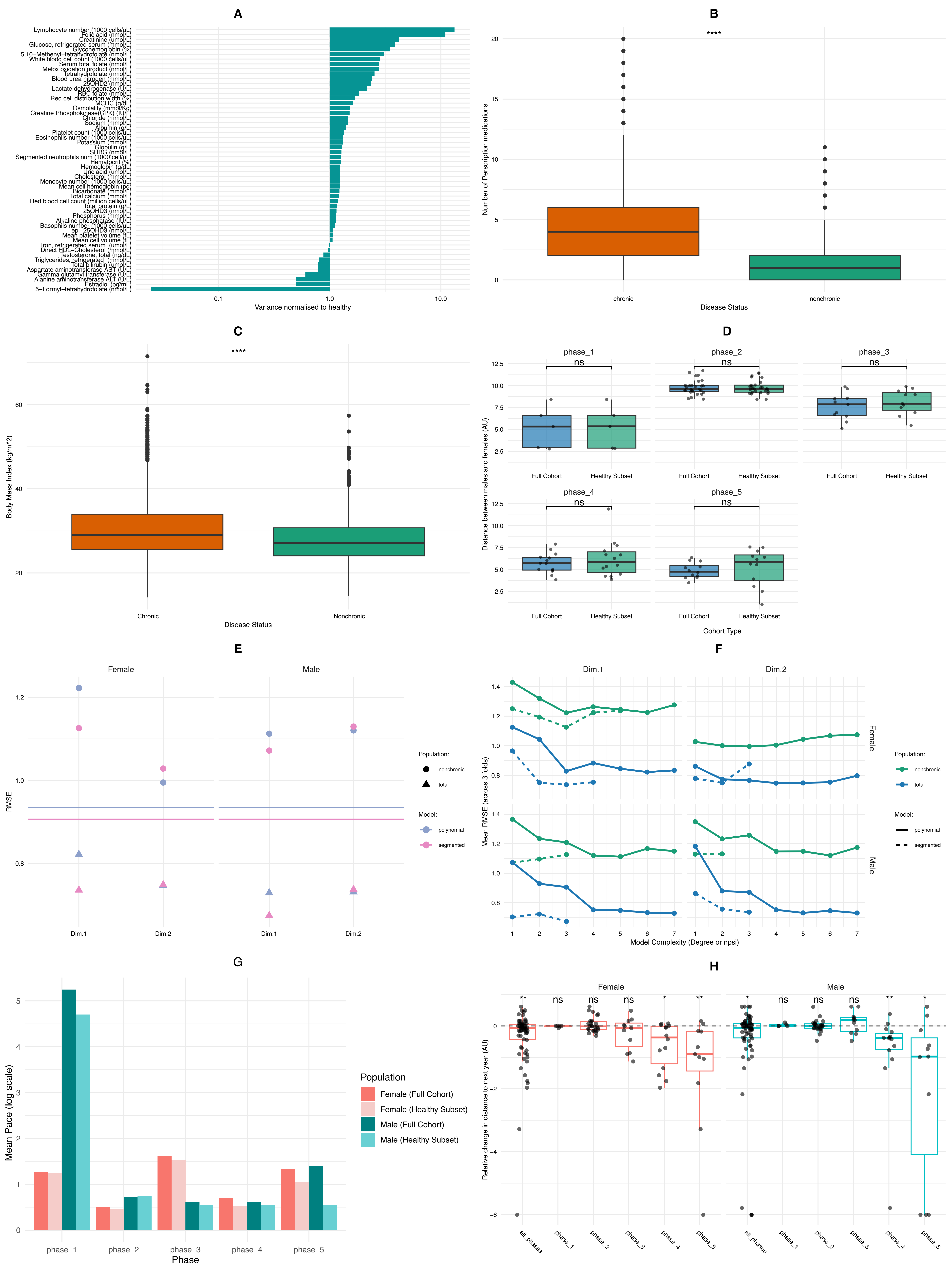

### Supplementary Figure 3

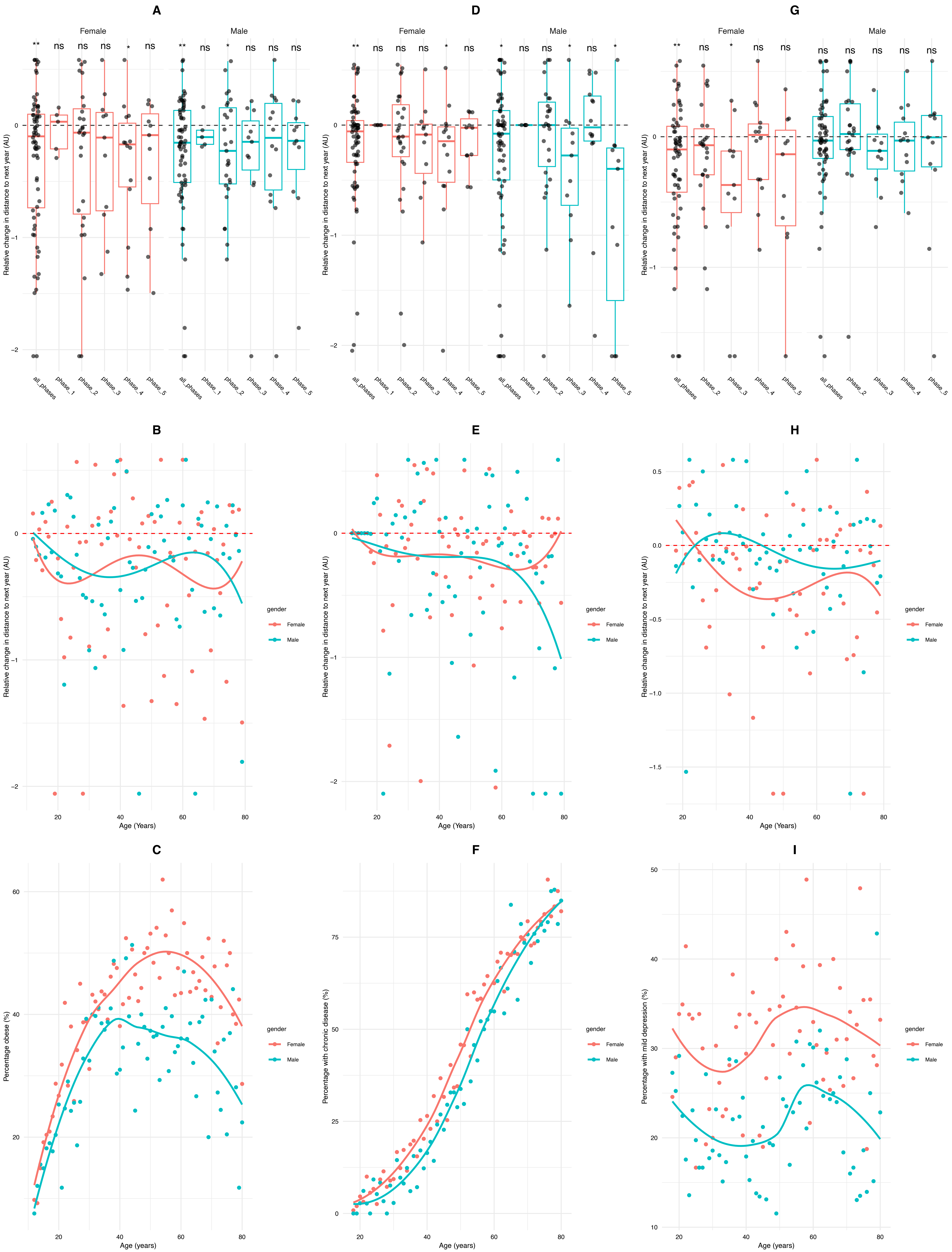
